# 3D Exploration of the Brainstem in 50-micron Resolution MRI

**DOI:** 10.1101/768002

**Authors:** N Makris, RJ Rushmore, P Wilson-Braun, G Papadimitriou, I Ng, Y N Rathi, F Zhang, LJ O’Donnell, M Kubicki, E Yeterian, JJ Lemaire, E Calabrese, GA Johnson, R Kikinis

## Abstract

The brainstem, a structure of vital importance in the mammals, is currently becoming a principal focus in cognitive, affective and clinical neuroscience. Midbrain, pontine and medullar structures are the epicenter of conduit, cranial nerve and such integrative functions as consciousness, emotional processing, pain and motivation. In this study, we parcellated the nuclear masses and the principal fiber pathways that were visible in a high resolution T2-weighted MRI dataset of 50-micron isotropic voxels of a postmortem human brainstem. Based on this analysis, we generated a detailed map of the human brainstem. To assess the validity of our maps, we compared our observations with histological maps of traditional human brainstem atlases. Moreover, we reconstructed the motor, sensory and integrative neural systems of the brainstem and rendered them in 3D representations. We anticipate the utilization of these maps by the neuroimaging community at large for applications in basic neuroscience as well as in neurology, psychiatry and neurosurgery, due to their versatile computational nature in 2D and 3D representations in a publicly available capacity.

## Introduction

The human brainstem, a structure the size of a thumb yet of the utmost biological importance, remains one of the most inaccessible parts of the brain in basic and clinical neuroscience. Phylogenetically, the brainstem is the oldest part of the brain. It has three main structural divisions, namely the midbrain (or mesencephalon), pons (Latin for “bridge”) and medulla oblongata (“oblong core” as well as “bulbus” in Latin) rostro-caudally (Carpenter & Sutin, 1983) (Nieuwenhuys et al., 2008). Located in the posterior cranial fossa between the spinal cord and the diencephalon, it develops between the fifth and seventh week of embryonic life in humans (Nolte et al., 2016), and is strategically positioned to serve as a conduit in processes associated with ascending and descending fiber tracts as well as for cranial nerve-related and higher integrative functions (Carpenter & Sutin, 1983) (Nieuwenhuys et al., 2008) (Nolte et al., 2016). These functions are reflected by the fine morphological, topographical and qualitative structural architecture of the brainstem, namely the cellular composition of the individual nuclei and their fiber connections. From a neural systems perspective, specialized functions of the brainstem are more clearly understood with respect to sensory and motor systems for which the midbrain, pons and medulla oblongata are key structures, serving most cranial nerve nuclei. By contrast, the structural and functional anatomy of brainstem systems integrating consciousness, pain and different types of affect is less clear. Brainstem cytoarchitecture and subsequently its myeloarchitecture have been described to a considerable extent since the early 1900s in classical studies that paved the way for more recent mapping of its nuclear masses and fiber connections (Olszewski & Baxter, 1982) (Swanson, 2015). The atlases resulting from such investigations have enabled us to determine the location and cellular composition of individual nuclei as well as their fiber pathways in the human brainstem. The latter have been crucial in understanding normative neuroanatomy as well as neuropathology (Olszewski & Baxter, 1982).

With the advent of MRI, pioneering studies of the brainstem have demonstrated the potential of neuroimaging for analyzing this brain component structurally, metabolically and functionally (Toga & Mazziotta, 2002) (Sclocco et al., 2018). Currently, ex-vivo acquisitions of human brain tissue using high field MRI scanners have allowed for high spatial resolution brainstem datasets. These datasets have enabled the visualization of gray and white matter structures to such a degree that the goal of matching histological preparations traditionally used in neuroanatomy with those of the MRI images has been met for certain nuclei as a whole although not yet at the cellular level. Precise MRI-based 2D and 3D computational reconstructions of gray and white matter anatomical structures allows their integration into current multimodal neuroimaging. Consequently, these data can be used in the different fields of basic and clinical neuroscience in which knowledge of structure, function and metabolism of the human brainstem is needed.

In this study, we parcellated the anatomical gray matter structures and the fiber pathways of conduit and cranial nerve systems that were visible in an ultrahigh resolution T2-weighted MRI dataset of 50-micron isotropic voxels of a postmortem human brainstem. Based on our initial analyses, we combined the gray and white matter parcellation results to generate a detailed map of the human brainstem. To evaluate the anatomical accuracy and precision of our maps, we compared our observations with histological maps of classical human brainstem atlases. Furthermore, by assembling the different component structures, we reconstructed the motor, sensory and integrative neural systems of the brainstem and rendered them in 3D representations. Given the computational nature of our maps in 2D and 3D spaces, we anticipate their utilization by the neuroimaging community at large for applications in basic neuroscience as well as in neurology, psychiatry and neurosurgery.

## Methods

### Imaging acquisition and protocol parameters of an ultrahigh-resolution ex-vivo T2-weighted MRI dataset

Postmortem imaging was performed in a 7 Tesla small animal MRI system by Calabrese and colleagues (2015) as follows. RF transmission and reception were achieved with a 65 mm inner-diameter quadrature RF coil. Anatomical images were acquired using a 3D gradient echo pulse sequence with repetition time (TR) = 50 ms, echo time (TE) = 10 ms, flip angle (a) = 60 degrees, and bandwidth (BW) = 78 Hz/pixel. The field of view (FOV) was 80 × 55 × 45 mm, and the acquisition matrix was 1600 × 1100 × 900 resulting in a 50-micrometer isotropic voxel size. Total acquisition time was 14 hours (Calabrese, et al., 2015).

### Anatomical image analysis: Segmentation and labeling of regions of interest

Identification and delineation of anatomical regions of interest (ROI) was done on the T2-weighted MRI dataset as follows. First, we down-sampled the T2 MRI dataset using 3D Slicer; as a result of this operation, the original dataset consisting of 50-micrometer thick axial sections resulted in axial sections of 250-micrometers thickness. We then used the 3D Slicer software analysis platform to perform segmentation and labeling of brainstem anatomical structures, which were visualized (Federov et al., 2012). All the tools used are publicly available and can be downloaded from the official 3D Slicer website (www.slicer.org). Segmentation of structures was done as follows. The brainstem was segmented as a whole and then subdivided into three component parts, namely the midbrain or mesencephalon, the pons and the medulla. Following traditional anatomical conventions, the limiting axial plane between the midbrain and the pons was set at the uppermost extent of the basis pontis, whereas the limiting axial plane between the pons and the medulla was set at the lowermost extent of the basis pontis (DaSilva, et al., 2002). This yielded a gross morphological definition and topographical delimitation of the brainstem in the context of traditional neuroanatomical descriptions. Subsequently, segmentation of gray matter and white matter structures was carried out individually for each nucleus and fiber tract to the extent that the data would allow visualizing and identifying the individual structures. This was done using the Editor module of 3D Slicer, which allows segmentation operations. Once a structure was segmented it was labeled. Two expert neuroanatomists (RJR and NM) and two trained research assistants (PWB and IN) were involved in this process. To ensure anatomical accuracy in the identification of gray and white matter structures we were also guided by anatomical textbooks and atlases of the brainstem. These were Olszewski-Baxter (Olszewski & Baxter, 1982), Paxinos and Huang (Paxinos & Huang, 1995), Haines (Haines, 1991), Carpenter and Sutin, (Carpenter & Sutin, 1983), and Nolte (Nolte et al., 2016; Vanderah, 2018). To address neural systems analysis in the brainstem, we assembled individual structures into motor, sensory and integrative neural systems, a topic of great relevance in current basic and clinical neuroscience (Panksepp, 2004) (Mesulam, 2000) (Damasio, 2010) (Solms & Turnbull, 2002).

## Results

Gross segmentation of the brainstem and its three principal parts, namely midbrain, pons and medulla, was readily accomplished. At an individual structure level of analysis, we were able to identify with confidence forty-seven (47) gray and white matter structures within the entire brainstem and the cerebral aqueduct. These consisted of twenty-five (25) gray matter structures, twenty-two (22) fiber tracts and the cerebral aqueduct. Of the forty-seven structures that were identified, we were able to delineate morphometrically all twenty-five gray matter structures, sixteen (16) white matter structures and the cerebral acqueduct as listed in detail in Table 1 and illustrated in Figure 1 in six representative axial sections. For anatomical reference and clarity, we displayed our results side-by-side with corresponding plates of the Nolte textbook (Nolte et al., 2016) (Vanderah, 2018). The latter was done for didactic purposes to aid the student of brainstem anatomy who is not sufficiently familiar with this complex structure. The results observed in our analysis matched the topography reported by classical atlases, namely those of Olszewski-Baxter (1982), Paxinos and Huang (1995), and Haines (1991). The size of our brainstem dataset (i.e., 54.2 mm in length) was comparable to the sample used by Olszewski and Baxter (1982) (i.e., 54.02 mm) by 99.7% as well as with the brainstem analyzed by Paxinos and Huang (1995) (i.e., 59 mm) by 91.6%. The topographical relationships among the different gray and white matter structures and the ventricular complex also met expectations based on traditional neuroanatomy. At another level of analysis, the neural systems associated with brainstem structures were delineated and reconstructed in three-dimensional space as shown in Figure 2. These were the motor and sensory systems of the face (Fig. 2A), the eye motor system (Fig. 2B), the auditory system (Fig. 2C), the vestibular system (Fig. 2D), the vestibuloocular system (2E), the cerebellar systems (Fig. 2F), the motor and sensory body systems (Fig. 2G1 and 2G2) and, the integrative systems (Fig. 2H).

**Table 1:**
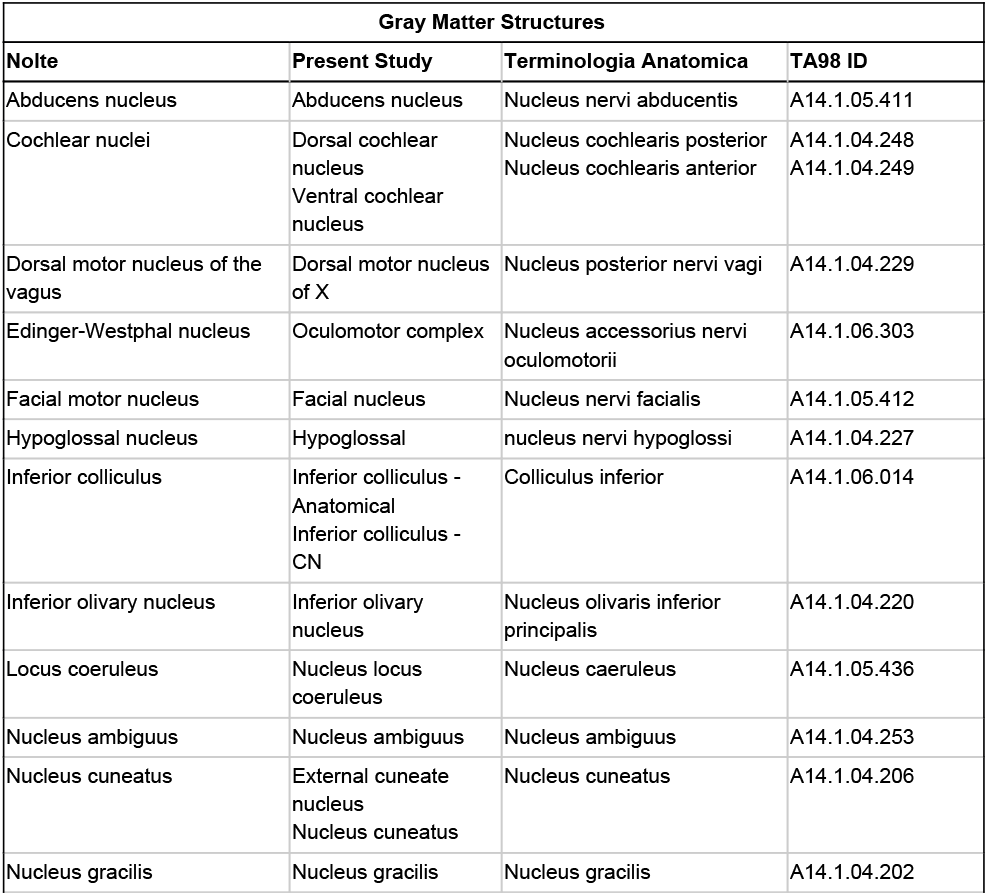
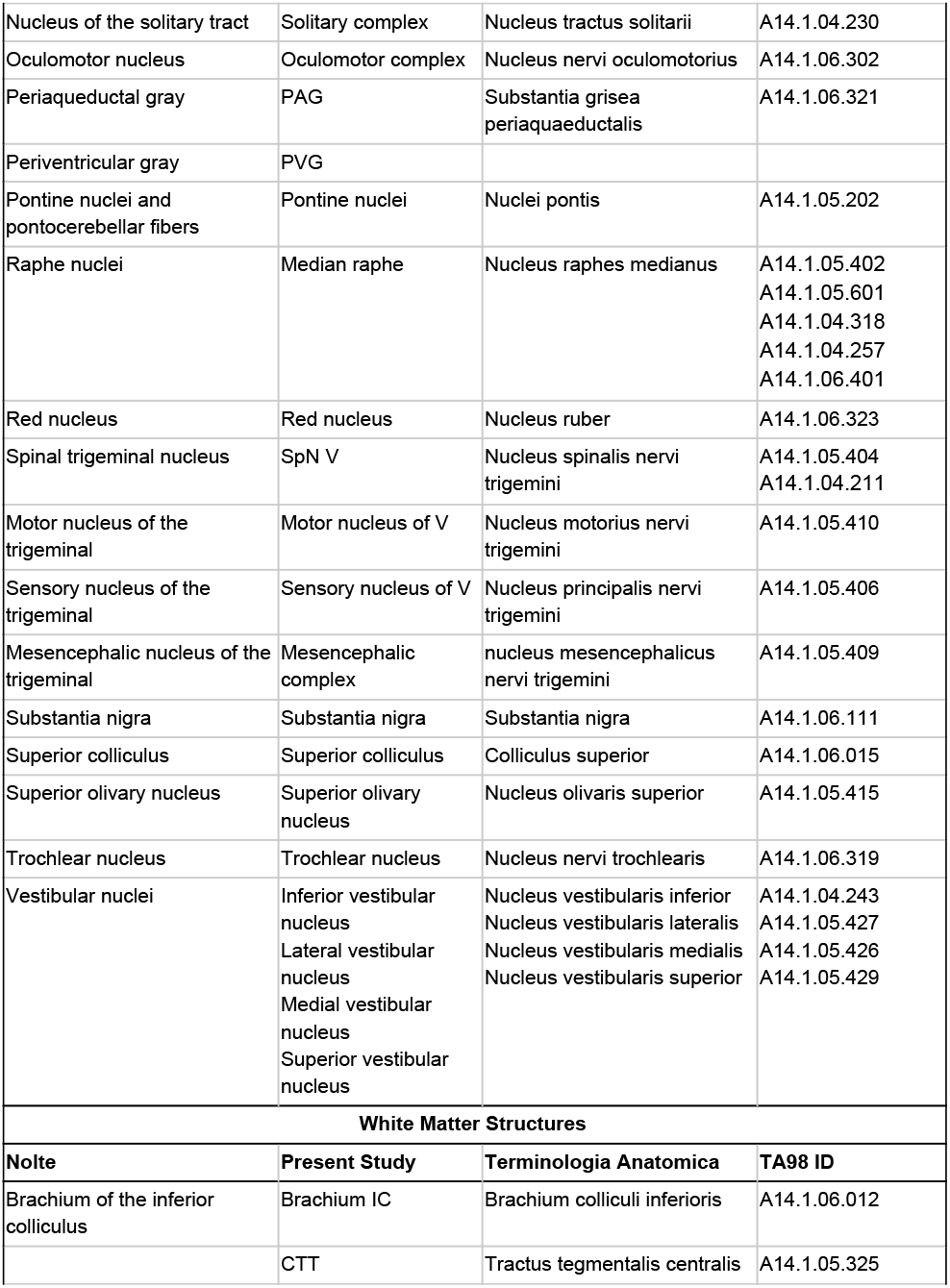
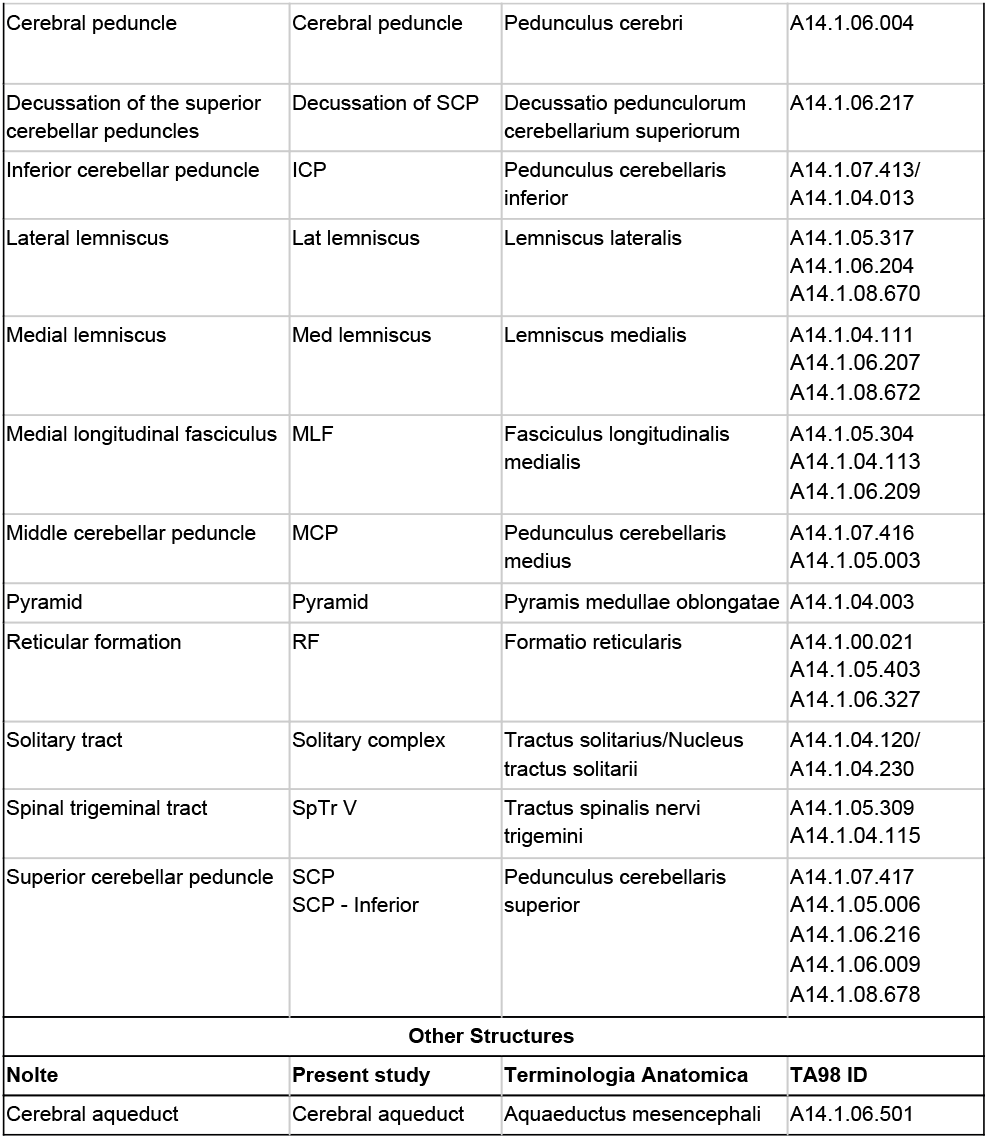
List of gray and white matter structures to relate brainstem structures as defined by Nolte (Nolte et al., 2016; Vanderah, 2018) with the present study and with the terminologia anatomica. The last column contains identification codes for the structures as found in the terminologia anatomica viewer (https://taviewer.openanatomy.org)

**Figure 1.**
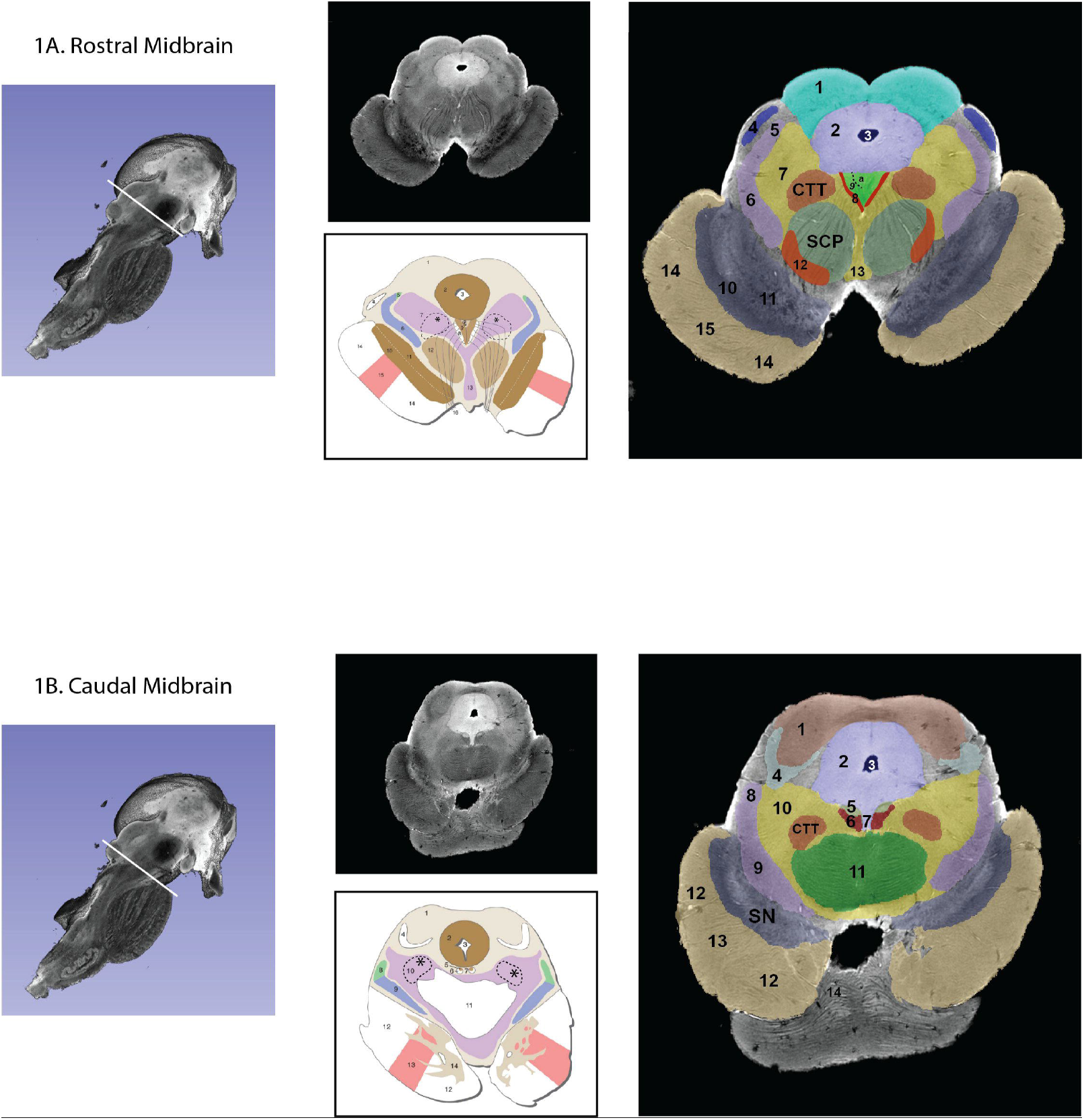
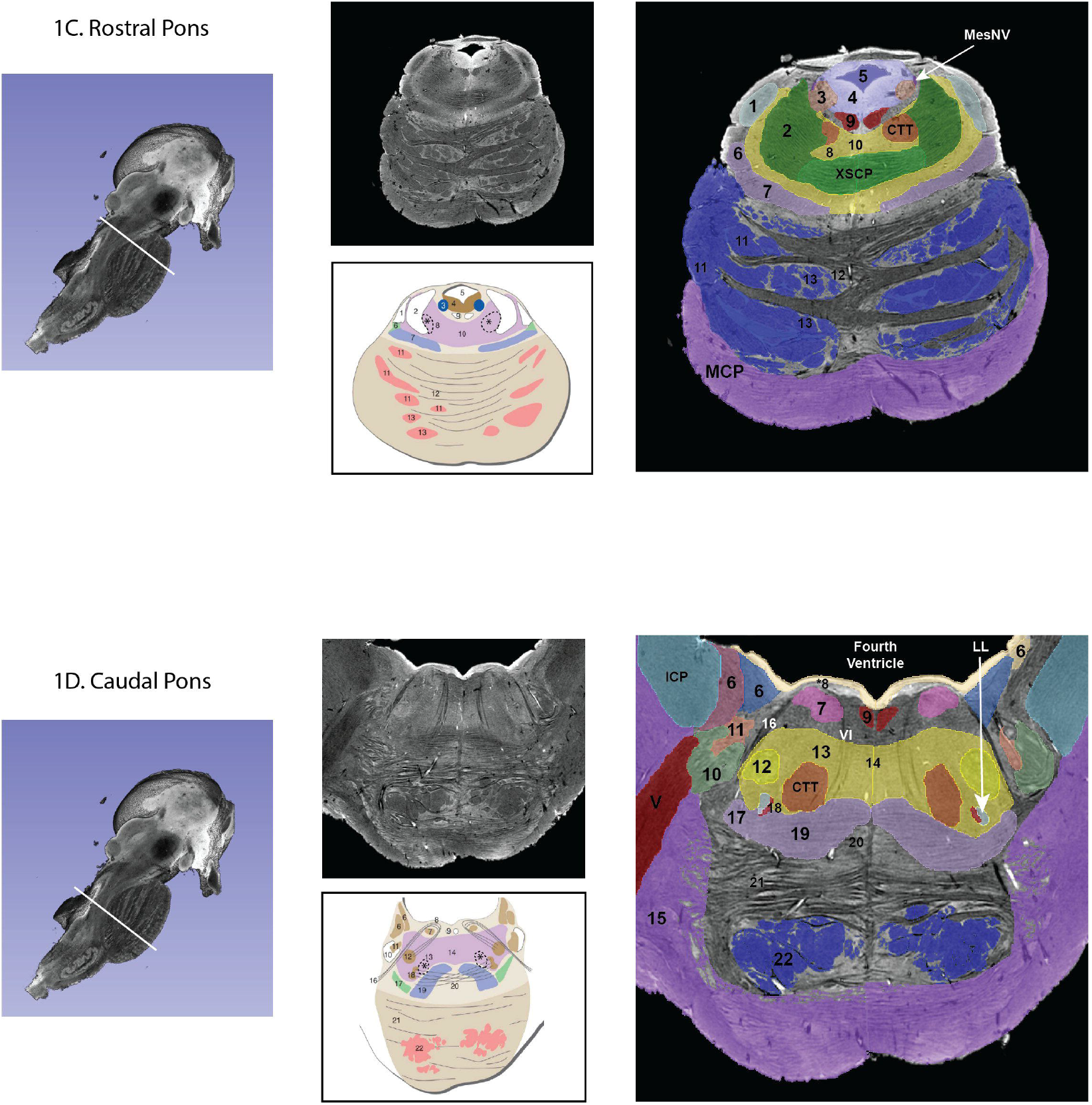
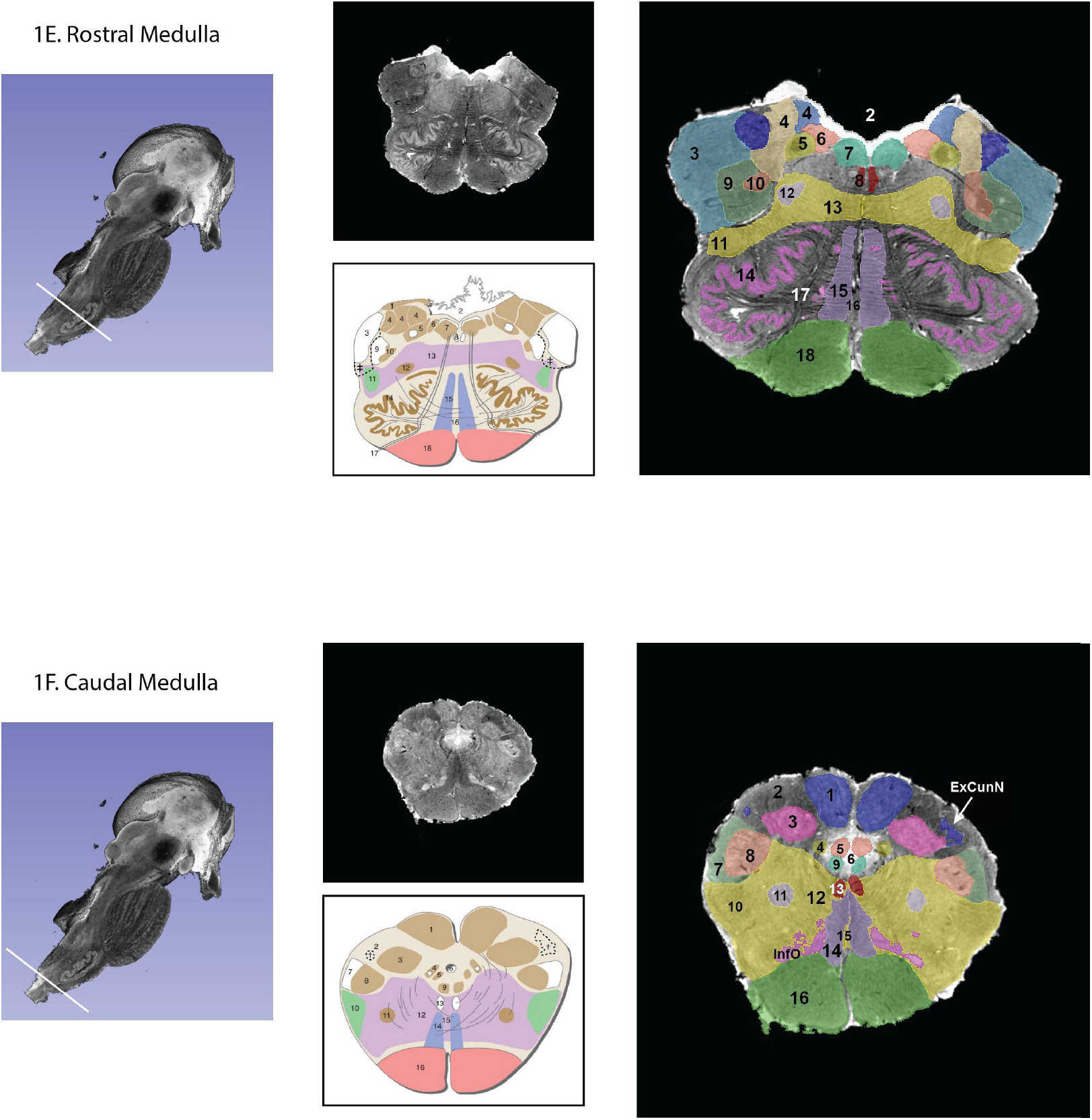
Transverse segmented sections through the midbrain (A, B), pons (C, D) and medulla (E, F) of the brainstem. For each subfigure, a hemisected longitudinal view of the brainstem is shown on the left with a white line representing the plane of section that corresponds to the images in the middle panel. The upper section of the middle panel is an image from the MRI dataset, and the lower picture is a schematic image from Essentials of the Human Brain (Vanderah, 2018) of a comparable brainstem section. The numbers in the schematic figures are recapitulated in the large segmented image on the right to show correspondence of structures between the MRI dataset and the schematic image. Note that 1) in some instances, some numbered structures present in the schematic figures do not appear in the segmented MRI image because they could not be reliably identified in the MRI image, and 2) some structures appear in the MRI image but not in the schematic images – these structures are identified by alphabetical abbreviation and reflect slight differences in the angle of cut or structures that were clear in the MRI but not demarcated in the schematic images. The red nucleus in particular, appears minimally in the MRI image due to the angle of section, which was taken at the caudalmost tip of the red nucleus. More specifically, this section represents the level where fibers of the superior cerebellar peduncle enter in the red nucleus. Furthermore, the central tegmental tract, which does not appear in the schematic figure has been identified in the MRI image. Moreover, fibers of the oculomotor nerve traversing the superior cerebellar peduncle are visible in both, the MRI image and the schematic figure. Abbreviations: Figure 1A 1. Superior Colliculus, 2. Periaqueductal gray, 3. Cerebral Aqueduct, 4. Brachium of the inferior colliculus, 5. Anterolateral system, 6. Medial lemniscus, 7. Reticular formation, 8. Medial longitudinal fasciculus, 9. Oculomotor nuclear complex (a. Nucleus of Edinger-Westphal), 10. Substantia Nigra pars reticulata, 11. Substantia nigra pars compacta, 12. Red nucleus, 13. Ventral tegmental area, 14. Corticopontine fibers of the Cerebral peduncle, 15. Corticospinal and corticobulbar fibers of the Cerebral peduncle. CTT – Central tegmental tract, SCP – Superior cerebellar peduncle. The CTT was added in the schematic image from Essentials of the Human Brain (Vanderah, 2018) in a dashed line indicated by an asterisk. Figure 1B 1. Inferior colliculus, 2. Periaqueductal gray, 3. Cerebral aqueduct, 4. Lateral lemniscus, 5. Trochlear nucleus, 6. Medial longitudinal fasciculus, 7. Raphe nuclei, 8. Anterolateral system, 9. Medial lemniscus, 10. Reticular formation, 11. Decussation of the superior cerebellar peduncle 12. Corticopontine fibers of the Cerebral peduncle, 13. Corticospinal and corticobulbar fibers of the Cerebral peduncle, 14. Pontine nuclei. CTT – Central tegmental tract, SN – Substantia nigra. The CTT was added in the schematic image from Essentials of the Human Brain (Vanderah, 2018) in a dashed line indicated by an asterisk. Figure 1C 1. Lateral lemniscus, 2. Superior cerebellar peduncle, 3. Locus coeruleus, 4. Periaqueductal gray, 5. Fourth ventricle, 6. Anterolateral system, 7. Medial lemniscus, 8. Reticular formation, 9. Medial longitudinal fasciculus, 10. Raphe nuclei, 11. Corticopontine fibers of the Cerebral peduncle, 12. Pontine nuclei, 13. Corticospinal and corticobulbar fibers. CTT – Central tegmental tract, MCP – Middle cerebellar peduncle, MesNV – Mesencephalic tract and nucleus of trigeminal, XSCP – Decussation of the superior cerebellar peduncle. The CTT was added in the schematic image from Essentials of the Human Brain (Vanderah, 2018) in a dashed line indicated by an asterisk. Figure 1D 6. Vestibular complex of nuclei, 7. Abducens nucleus, *8. The location of the internal genu of the facial nerve is indicated here, although it should be noted that in this MRI image the plane of section includes both, fibers of the facial nerve laterally (16 in this figure) and fibers of the abducens nerve medially (VI), 9. Medial longitudinal fasciculus, 10. Spinal tract of the trigeminal, 11. Spinal nucleus of the trigeminal, 12. Facial motor nucleus, 13. Reticular formation, 14. Raphe nuclei, 15. Middle cerebellar peduncle, 16. Facial nerve fibers, 17. Anterolateral system, 18. Superior olivary nucleus, 19. Medial lemniscus, 20. Trapezoid body, 21. Pontine nuclei, 22. – Corticospinal, corticobulbar and corticopontine fibers. CTT – Central tegmental nucleus, ICP – Inferior cerebellar peduncle, LL – Lateral lemniscus, MCP – Middle cerebellar peduncle, V – Trigeminal nerve fibers, VI – Abducens nerve fibers. The CTT was added in the schematic image from Essentials of the Human Brain (Vanderah, 2018) in a dashed line indicated by an asterisk. Figure 1E 2. Fourth Ventricle, 3. Inferior cerebellar peduncle, 4. Vestibular nuclei, 5. Solitary tract and nucleus of solitary tract, 6. Dorsal motor nucleus of the vagus, 7. Hypoglossal nucleus, 8. Medial longitudinal fasciculus, 9. Spinal trigeminal tract, 10. Spinal trigeminal nucleus, 11. Anterolateral system, 12. Nucleus ambiguous, 13. Reticular formation, 14. Inferior olivary nucleus, 15. Medial lemniscus, 16. Raphe nuclei, 17. Hypoglossal nerve fibers, 18. Pyramid. Please note that in the MRI image the cochlear nuclei are not present due to the plane of section. The border of the inferior cerebellar peduncle was extended in the schematic image from Essentials of the Human Brain (Vanderah, 2018) in a dashed line indicated by a double cross. Figure 1F. 1. Nucleus gracilis, 2. Fasciculus cuneatus, 3. Nucleus cuneatus, 4. Solitary tract and nucleus of solitary tract, 5. Dorsal motor nucleus of the vagus, 7. Spinal trigeminal tract, 8. Spinal trigeminal nucleus, 9. Hypoglossal nucleus, 10. Anterolateral system, 11. Nucleus ambiguous, 12. Reticular formation, 13. Medial longitudinal fasciculus, 14. Medial lemniscus, 15. Raphe nuclei, 16. Pyramid, InfO – Inferior olivary nucleus. The external cuneate nucleus (ExCunN) was added in the schematic image from Essentials of the Human Brain (Vanderah, 2018) in a dashed line indicated by a single cross.

**Figure 2.**
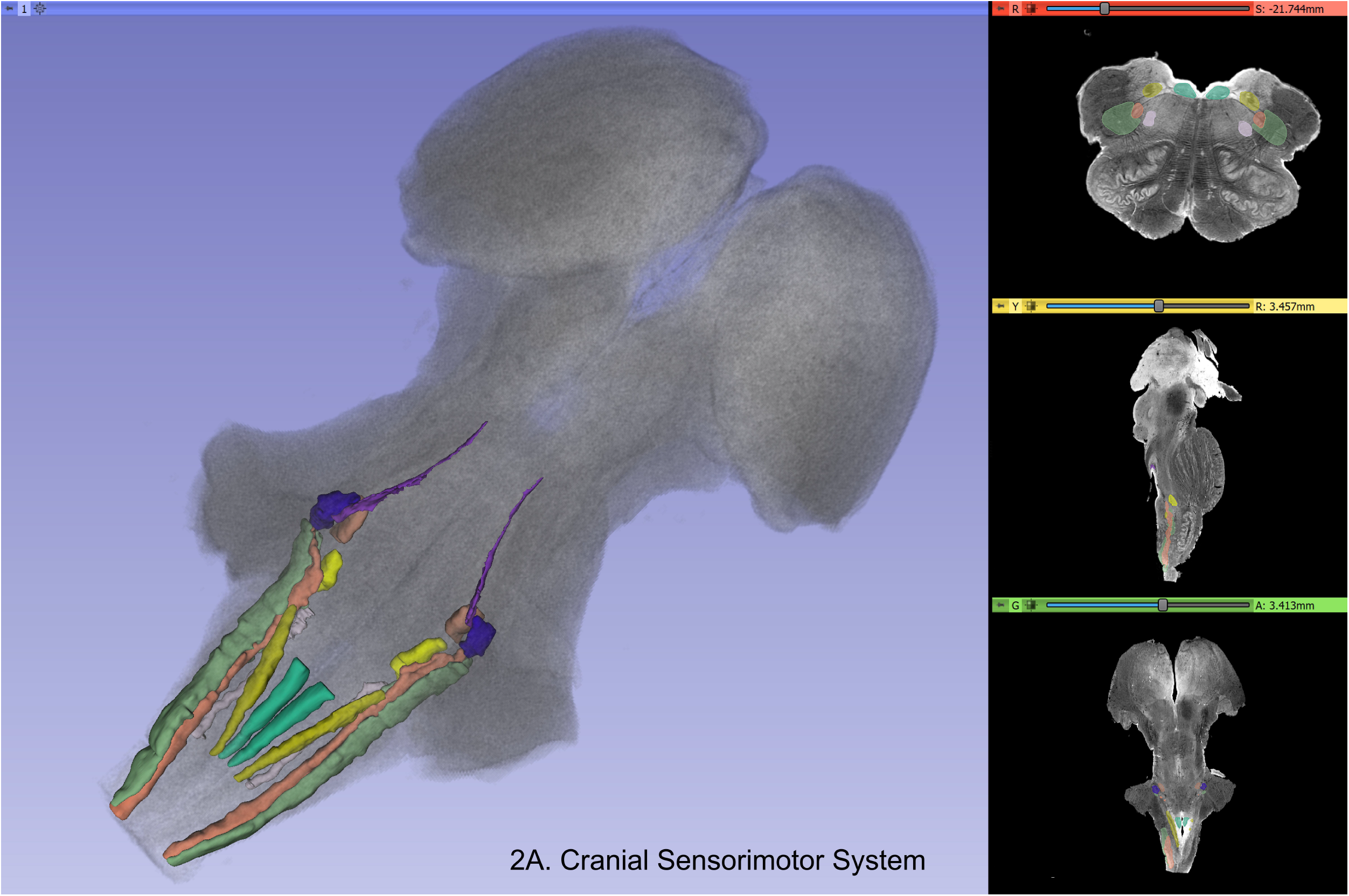
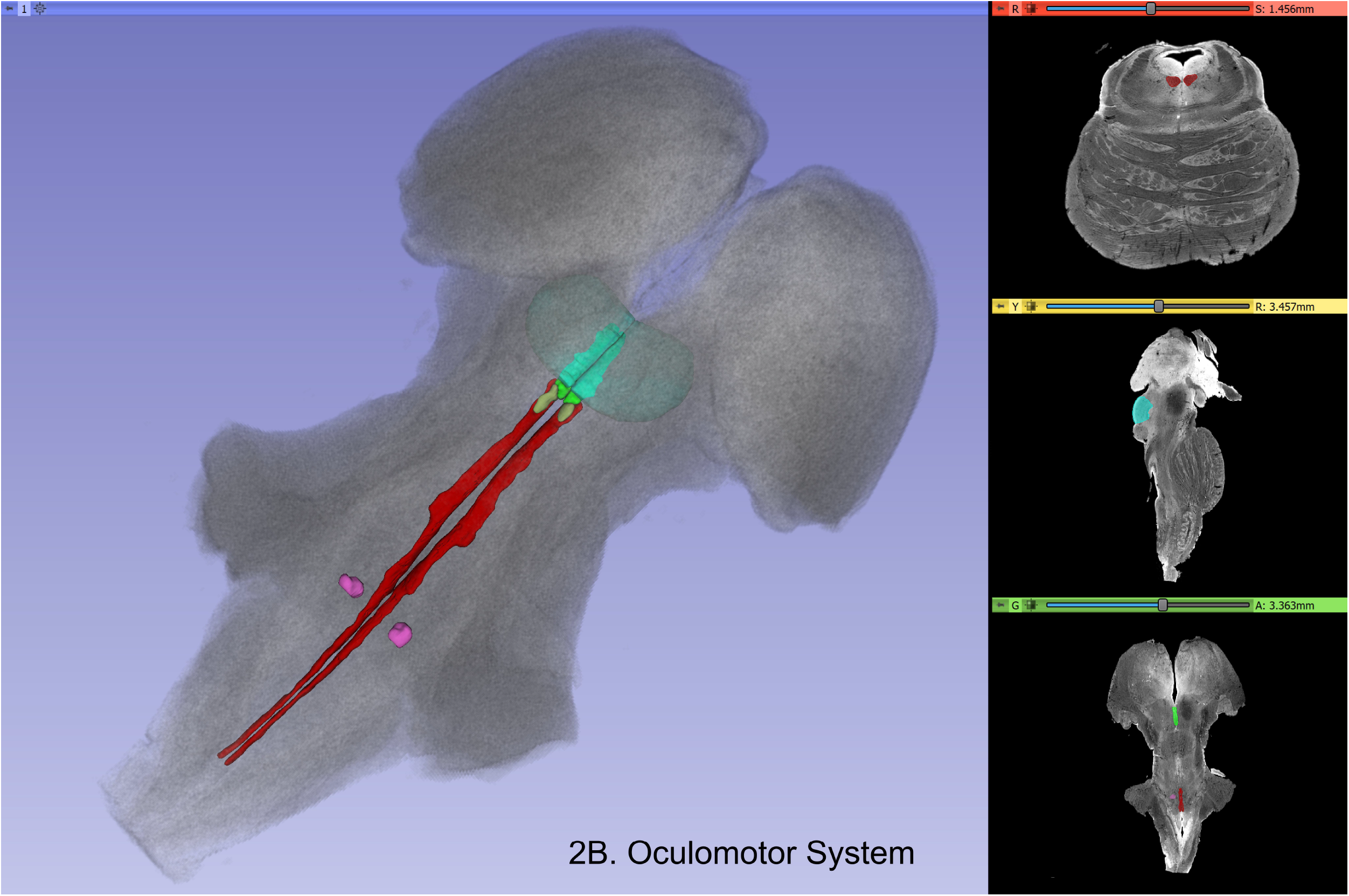
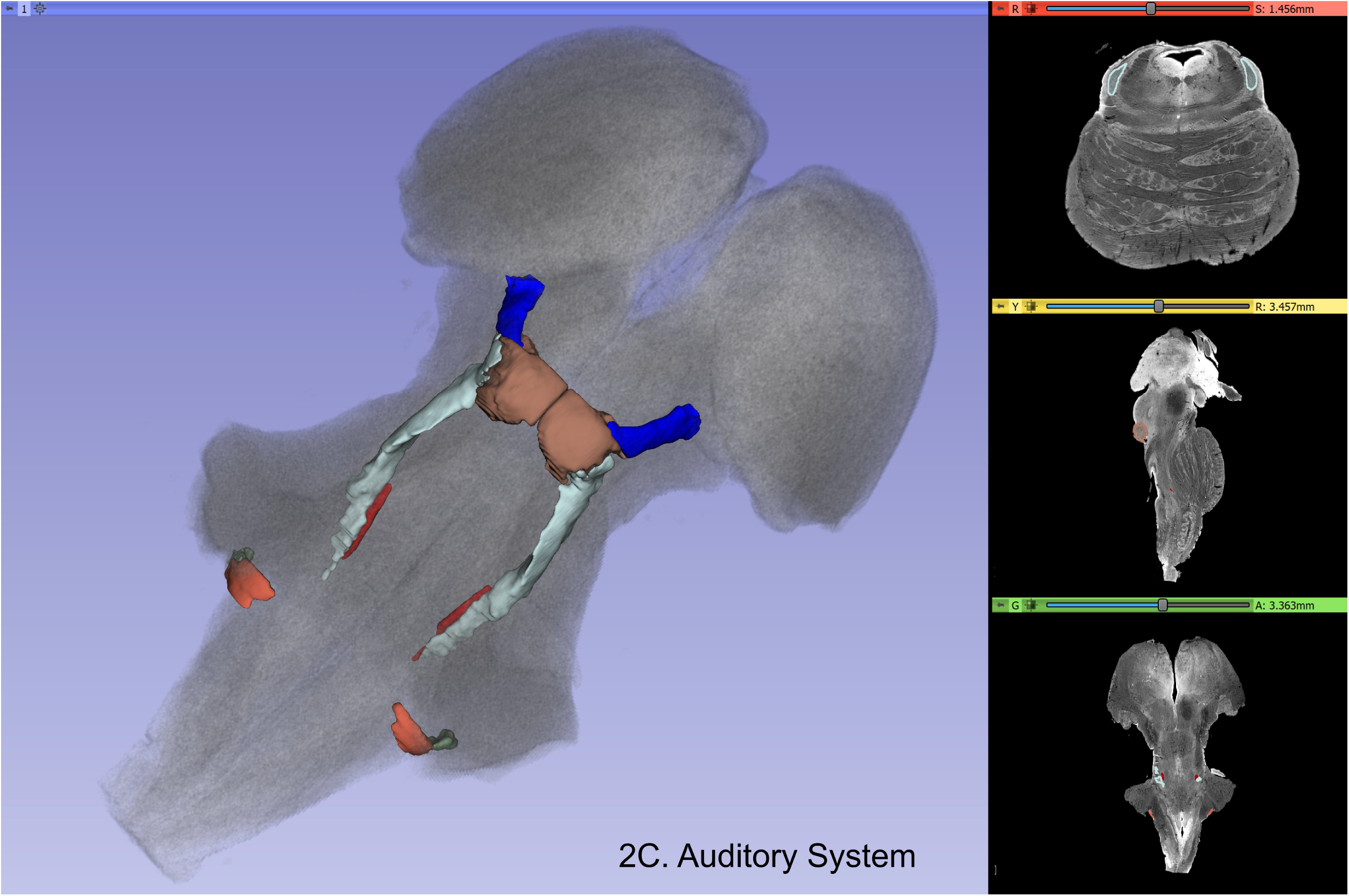
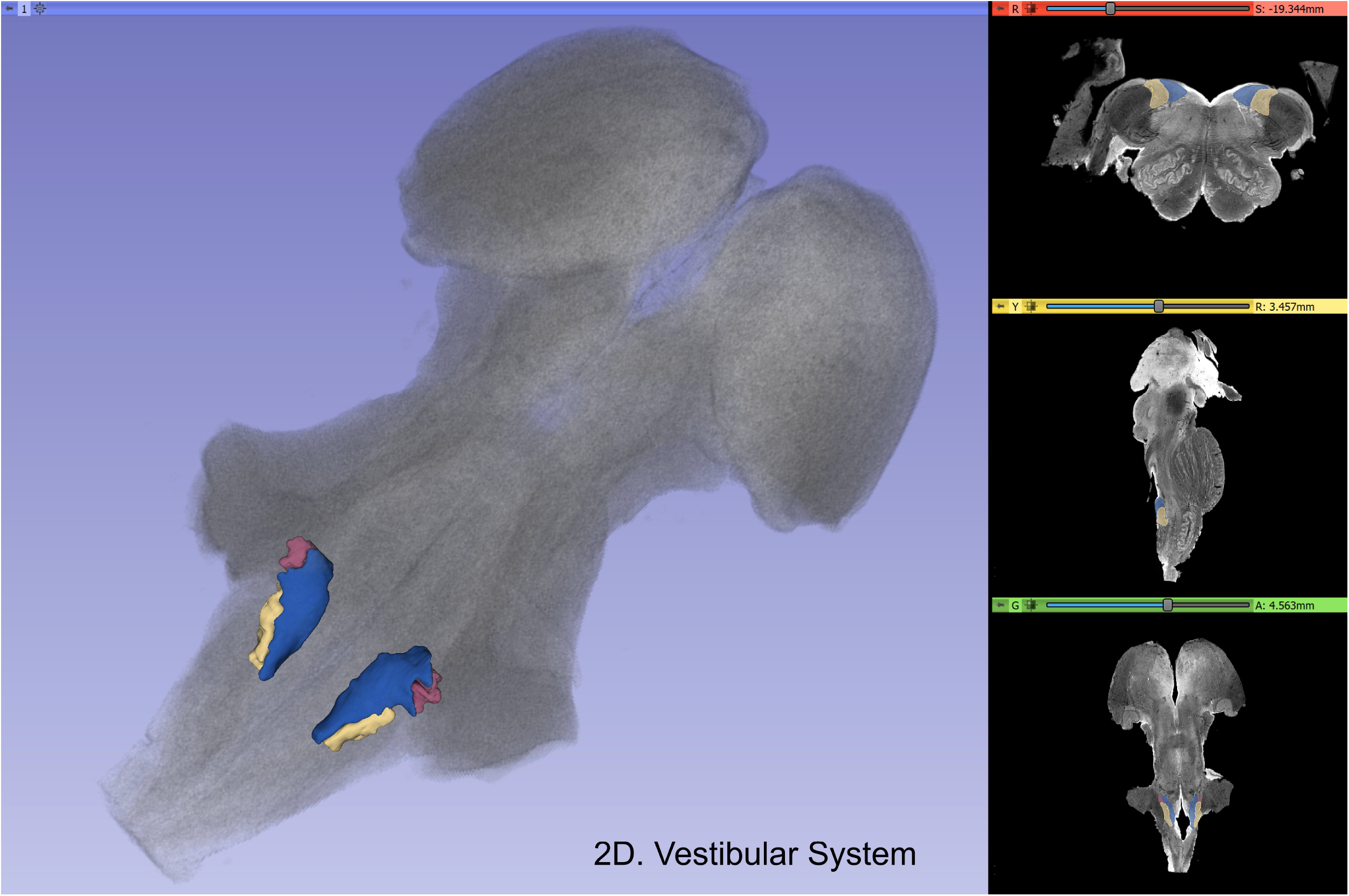
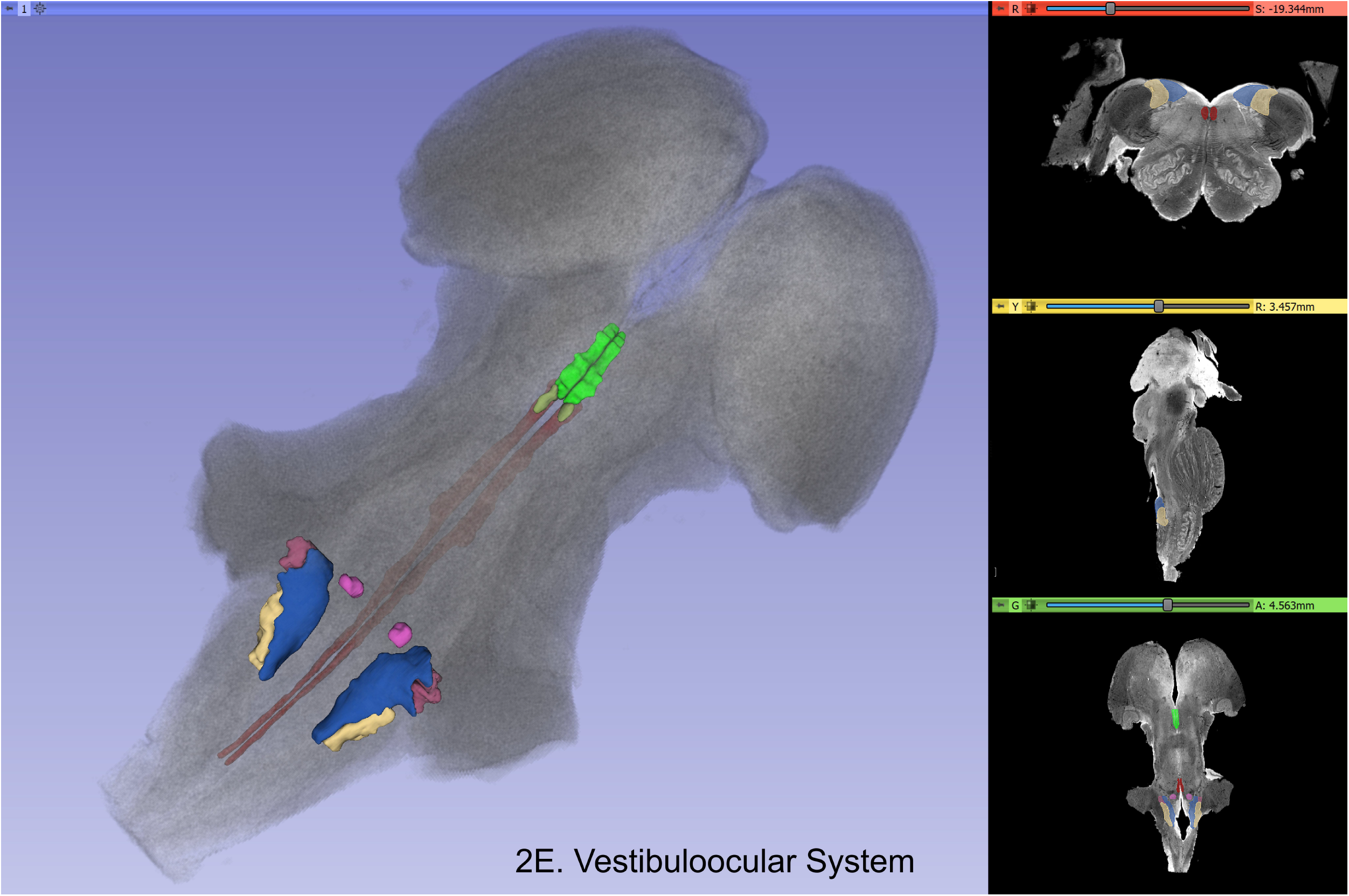
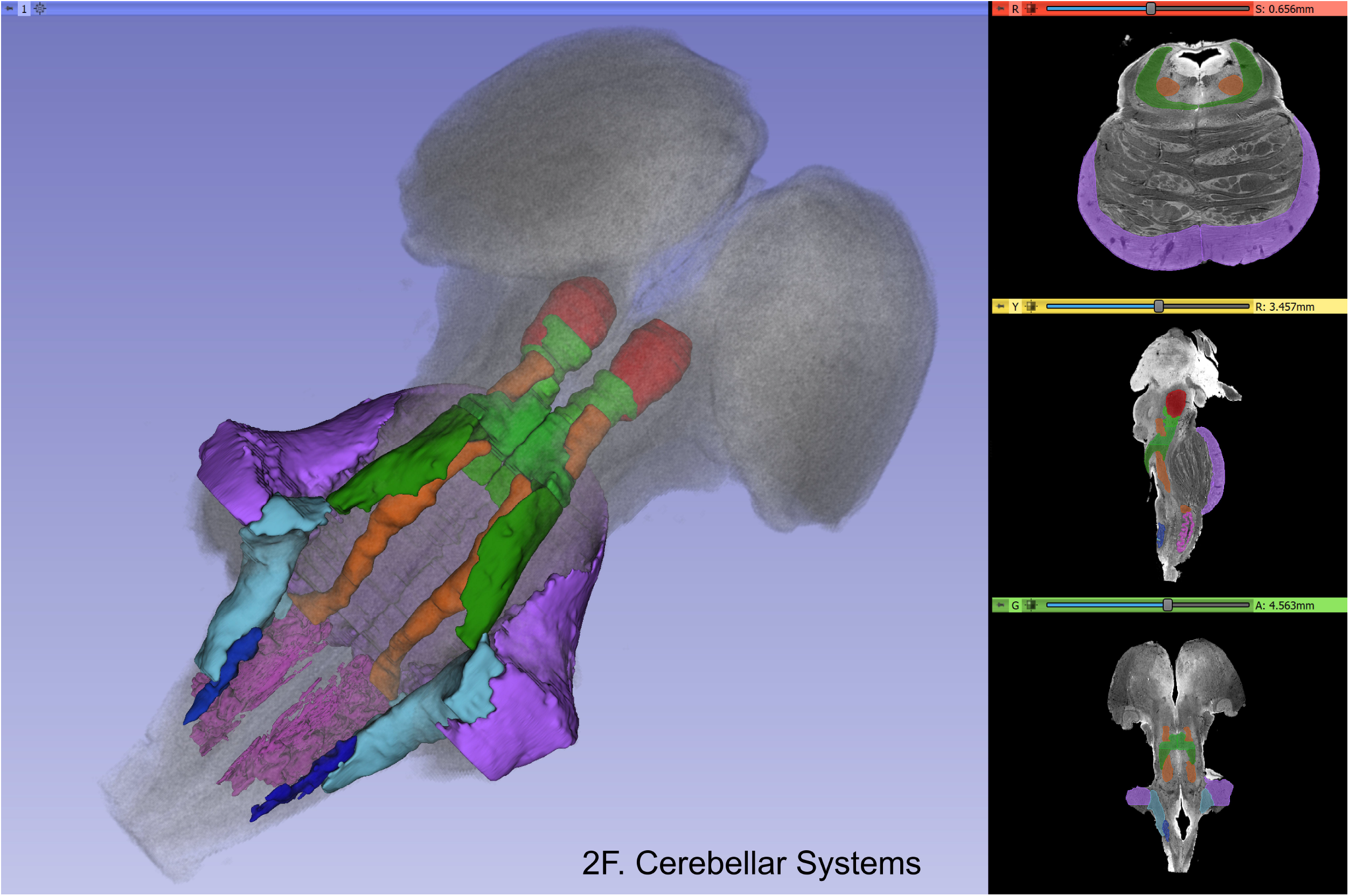
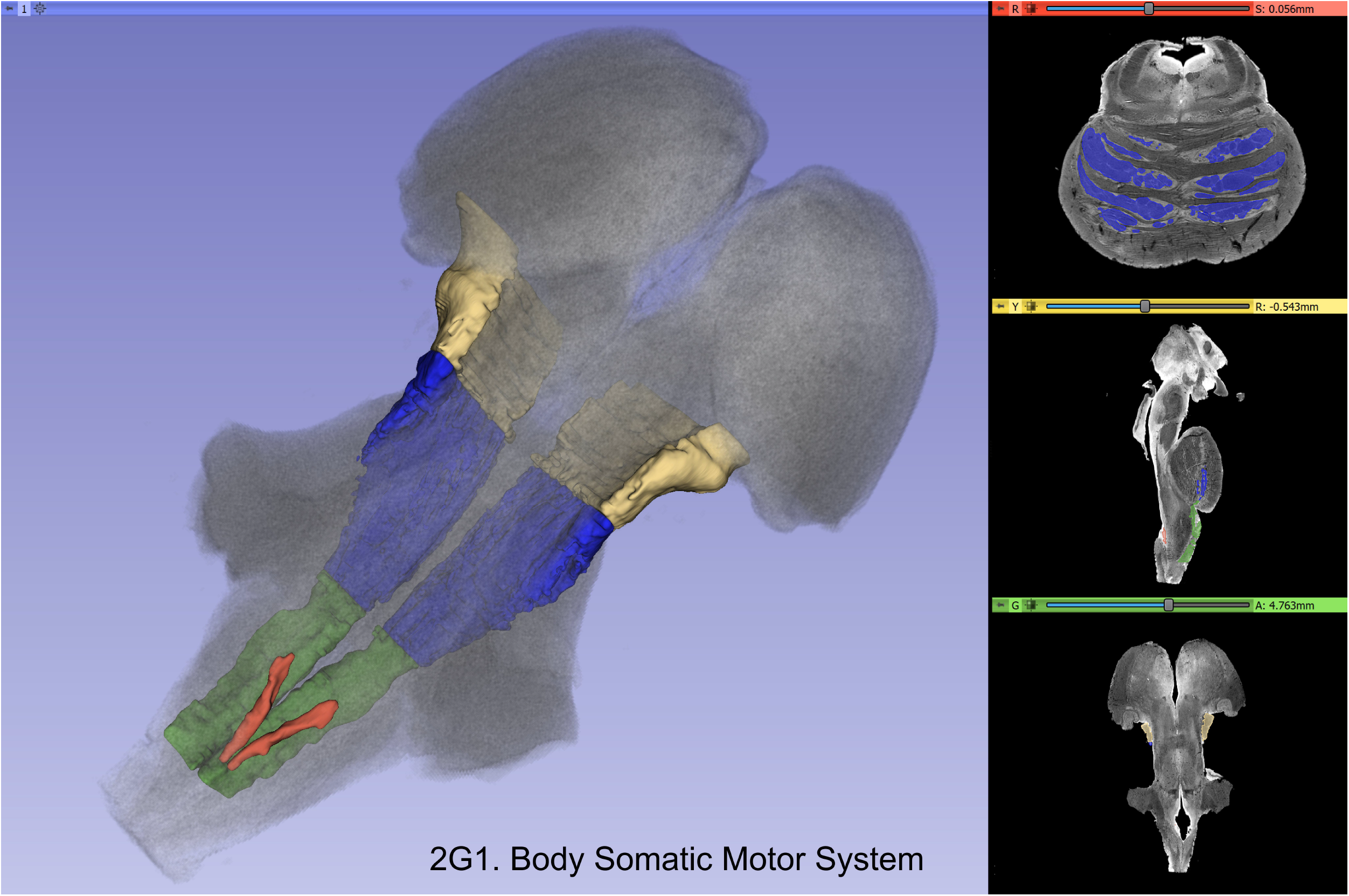
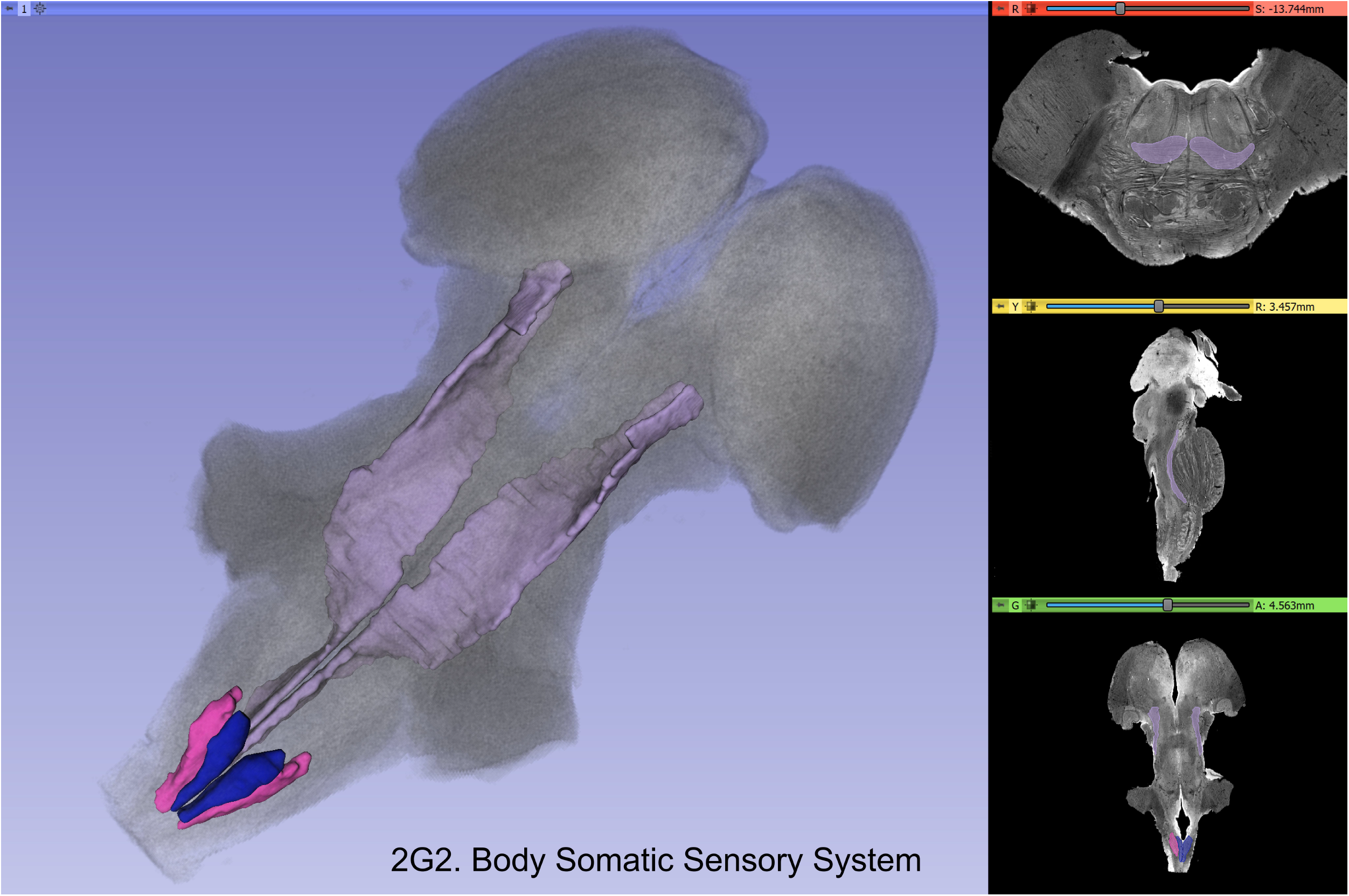
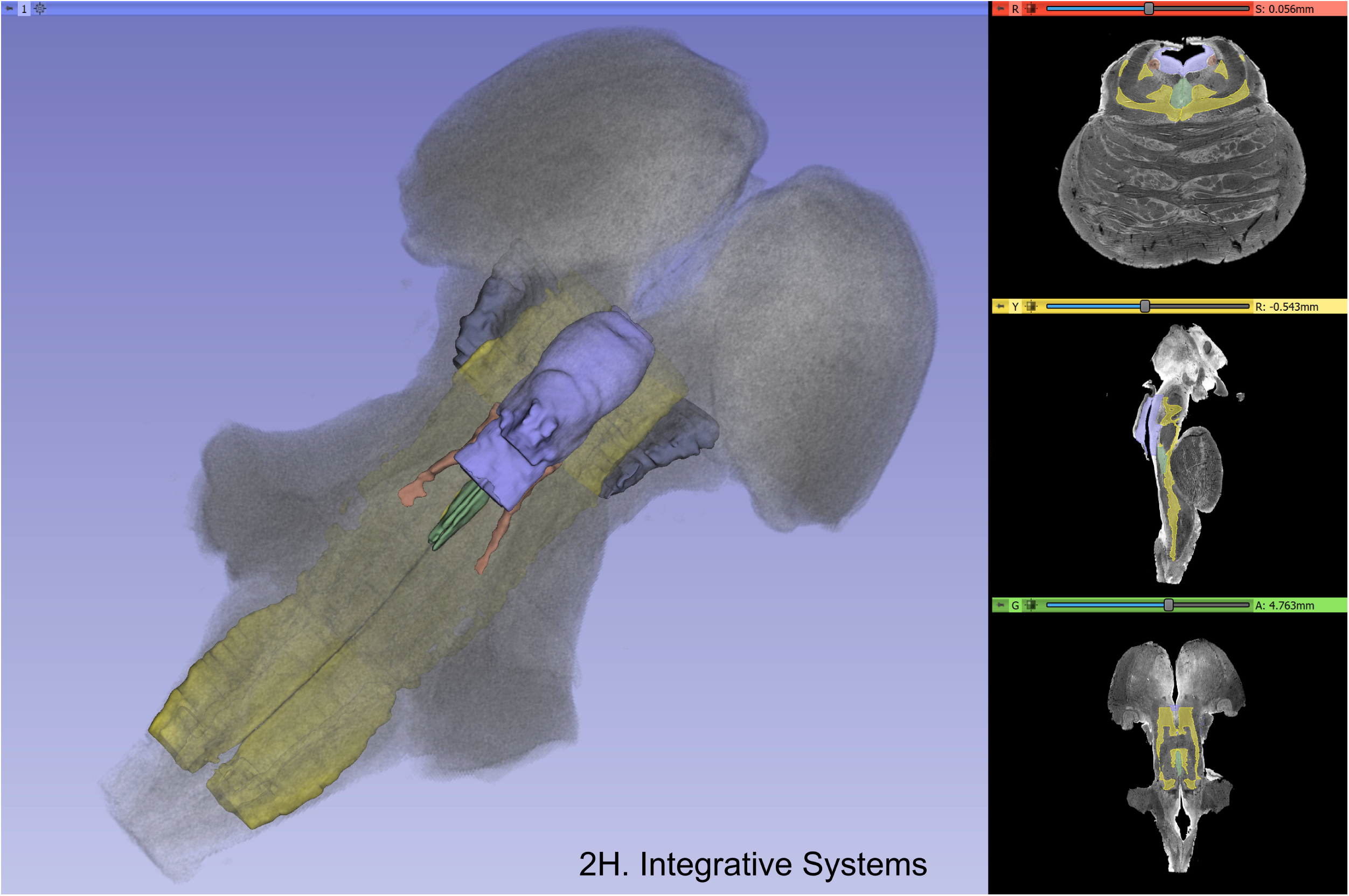
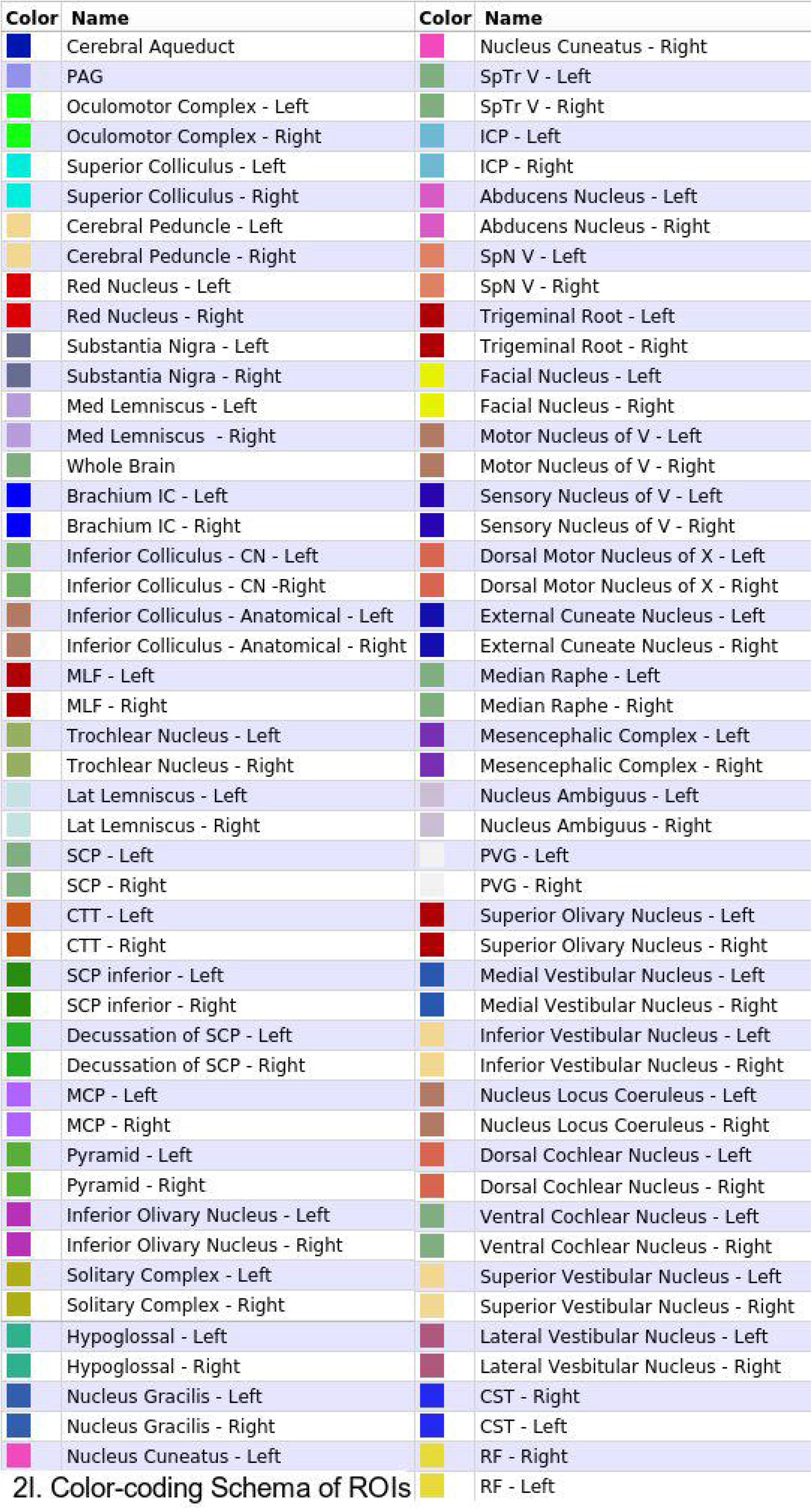
Three dimensional reconstructions of select brainstem systems. For each subfigure, the three-dimensional rendering is overlaid on a volume rendering of the brainstem, and selected slices in the axial (red), coronal (yellow), and sagittal (green) planes are illustrated on the right. To generate the three-dimensional structures, a 0.5mm median smooth was applied. A. Sensory and motor cranial system, B. Oculomotor system, C. Auditory system, D. Vestibular system, E. Vestibuloocular system, F. Cerebellar systems, G. Motor (G1) and Sensory (G2) systems of the body, H. Integrative systems, and I. Color-coding schema of regions of interest (ROIs). Abbreviations in the color-coding schema are explained in Table 1. Note that the most clearly resolvable structures on the MRI images were segmented and reconstructed and that these images do not encapsulate all component structures of each system.

## Discussion

In this study, we localized and delineated to a considerable extent the principal structures of the human brainstem using an MRI ex vivo ultrahigh-resolution dataset with properties largely comparable to histological representation. More specifically, we volumetrically identified forty-seven gray (viz., twenty-five) and white matter (viz., twenty-two) structures in the brainstem based on ex-vivo human T2-weighted MRI data of 50-micron isotropic spatial resolution. Furthermore, we were able to delineate morphometrically all twenty-fine gray matter structures and sixteen of the twenty-two white matter structures, which were originally identified by JR and NM. To our knowledge, this has not been previously achieved in MRI-based morphometric analysis. Moreover, we assembled individual structures into neural systems specifically carrying-out cranial nerve, conduit and integrative functions.

The brainstem is becoming a major focus in clinical and basic neuroscience due to our current ability to image it volumetrically at resolutions that were previously inaccessible to MRI. Although relatively small in size, approximately the size and shape of a thumb in humans, this brain structure is associated with vital biological integrative functions such as consciousness, motivation, pain and reward as well as cranial nerve biology and conduit somatosensory and motor functions. Thus, the brainstem is critical for a complete understanding of brain structure-function relationships. We expect our anatomical analysis to be of use to neuroanatomists, neuropathologists, neurologists as well as psychiatrists and basic neuroscientists.

Early students of neuroanatomy who macro-dissected the human brain provided gross, approximate descriptions of the brainstem as a whole and its component parts, namely the midbrain, pons and medulla (Vesalius, 1543) (Varoli, 1573) (Diemerbroeck, 1689) (Haller, 1747) (Baer, 1837). Microscopic studies of the human brainstem have provided us with comprehensive descriptions of this structure since the early 1900s. A historical review of these aspects of the human brainstem has been provided by Olszewski and Baxter (1982) and can be highlighted briefly as follows. Jacobshon’s drawings in particular (1909) have served as a classical guideline for anatomical work regarding the brainstem nuclei given their remarkable anatomical accuracy (Jacobsohn-Lask, 1909). The classical work of Ziehen has provided extensive descriptions of cyto- and myelo-architecture of fiber pathways in the brainstem (Ziehen, 1933). Further studies performed on brainstem cytoarchitectonics have provided additional detail and visualization capabilities with the use of photomicrographs (Gagel & Bodechtel, 1930) (Stern, 1935) (Crosby & Woodburne, 1943) (Riley, 1943). More recently, the work of Olszewski and Baxter (1982) and of Paxinos and Huang (1995) stands out for their in-depth descriptions of brainstem nuclei cytoarchitecture and topography. It should be noted that the atlas of Paxinos and Huang (1995) provides unprecedented detail and coverage of not only nuclei but also fiber tracts in the human brainstem – almost twice the number of fiber pathways compared to other atlases. Given that in the present study we needed to simplify and adapt the brainstem nuclear and fiber tract analysis to the neuroimaging data under analysis, we used primarily the Olszewski and Baxter (1982) atlas as well as the atlas of Haines (1991), while the Paxinos and Huang (1995), atlas served as the final comparison reference and testbed for our analyses. Finally, for didactic purposes we have displayed the human brainstem gray and white matter structures in six classical axial planes, namely through the rostral and caudal midbrain, rostral and caudal pons, rostral and caudal medulla, as generally accepted in neuroanatomy following, for example, the Nolte textbook (Nolte, 1999) (Nolte & Angevine, 1995) (Nolte et al., 2016) (Vanderah, 2018).

In more recent years, with the advent of MRI, there have been several studies addressing the in vivo and non-invasive visualization of the gray and white matter of the human brainstem (Salamon, et al., 2005) (Kamali et al., 2009) (Yang et al., 2011) (Linnman et al., 2012) (Bianciardi, et al., 2015) (Meola et al., 2016) (Sclocco et al., 2018). MRI-based morphometry considered the morphological and volumetric characterization of the brainstem since the early 1990s (Filipek et al., 1994). Although the gross nature of these investigations precluded their addressing the fine architecture of the brainstem, these early studies indicated the great potential of neuroimaging to study this structure in vivo and non-invasively. As neuroimaging technology and methods of analysis evolved, brainstem anatomical analysis advanced using structural T1- and T2-weighted MRI as well as diffusion MRI (dMRI) and dMRI tractography. Pioneering morphometric studies with structural imaging measured the major components of the brainstem, namely the midbrain, pons and medulla, and were used to localize functional activation of specific cranial nerve nuclei in combination with task-specific fMRI acquisitions (DaSilva, et al., 2002). Furthermore, specific nuclear masses have been identified and labeled using T1- and T2-weighted MRI (Linnman et al., 2012) (Bianciardi, et al., 2015). Moreover, using dMRI tractography, several fiber tract connections have been identified and delineated in the brainstem. Although most studies have addressed principally major motor and sensory connections such as the corticospinal tract (Salamon, et al., 2005) (Meola et al., 2016), the cerebellar peduncles (Meola et al., 2016), the corticopontocerebellar pathways (Habas & Cabanis, 2007), and the medial and lateral lemniscus (Kamali et al., 2009) (Meola et al., 2016), there are also investigations focusing on finer connections such as the rubrospinal tract (Yang et al., 2011) Meola et al., 2016), spinothalamic tract, medial longitudinal fasciculus, dorsal longitudinal fasciculus (Meola et al., 2016) and central tegmental tract (Kamali et al., 2009) (Meola et al., 2016).

In the context of a body of growing knowledge in the anatomical brainstem imaging field, to our knowledge the present study takes a step forward in identifying and delineating the greatest number of nuclear structures provided to date. This was achieved principally because of the unprecedented resolution, i.e., 50-micrometer isotropic voxels, and the very high signal quality of the dataset we analyzed. Importantly, the anatomical analysis was done by expert neuroanatomists using 3D Slicer segmentation and visualization tools. Finally, comparisons of the delineated nuclear structures were done with guidance from classical brainstem textbooks and atlases portraying the precise cytoarchitecture and topography of the human brainstem nuclei. By assembling the component structural parts in the brainstem, we also have illustrated a viable means of visualizing in 3D the different cranial nerve systems, conduit systems and integrative systems, a most useful method in illustrating and understanding brainstem anatomy for basic neuroscientists as well as for neurologists, psychiatrists and neurosurgeons. As a result of this endeavor, 3D visualizations in the publicly available platform of 3D Slicer allow the student of the brainstem to use the atlas provided herein readily and relatively simply as a learning and teaching tool for this highly complex domain of neuroanatomy.

### Functional considerations

The organization of brainstem activities can be generally categorized in a didactic although simplistic manner as conduit, cranial nerve, and integrative functions. The topographic arrangement of the brainstem in relation to the spinal cord, the cerebellum and the cerebrum makes it a natural route of passage for the numerous fiber tracts interconnecting these structures, thus justifying its role as *conduit of fibers of passage*. In this study, we labeled twenty-one and delineated fifteen of these fiber pathways, as shown in Table 1 and Figure 2. It should, be noted, however, that fibers of passage may also course through the nuclei of the midbrain, pons and medulla. Although three of the *cranial nerves*, namely the olfactory (cranial nerve I), optic (cranial nerve II) and accessory (cranial nerve XI), do not project primarily or directly to the brainstem, the other nine are anatomically associated with the brainstem. Herein we were able to label the majority of cranial nerve nuclei, namely of cranial nerves III, IV, V, VI, VII, VIII, IX, X and XII in greater detail, as shown in Table 1 and Figure 2. As such, the main associated functions are sensory, i.e., facial and taste, hearing, equilibrium, visceral thoracic and abdominal, chemoceptive and baroceptive, as well as motor, i.e., eye movements, pupil and lens function, chewing, facial expression, swallowing, speech, visceral thoracic and abdominal as well as tongue movements. The most intriguing and less explored role of the brainstem remains the one related to its *integrative functions*. This is apparent from a behavioral point of view, given the complexity of such state-dependent functions as consciousness or emotion (Mesulam, 2000) (Solms & Turnbull, 2002) the brainstem is intricately involved in. Briefly, a number of integrative functions take place at the level of the midbrain, pons and medulla, dealing with regulation of consciousness (Moruzzi & Magoun, 1949) (Solms & Turnbull, 2002), visceral activity (Nieuwenhuys et al., 2008) (Nolte et al., 2016), stress response (Feldman & Saphier, 1985) (Feldman et al., 1995), pain (Linnman et al., 2012) (Menant et al., 2016), and complex motor and sensory processing (Carpenter & Sutin, 1983) (Nolte et al., 2016). A critical role in integrative functions is served by the reticular formation, as reflected structurally by the diffuse pattern of its connections with several other parts of the brain. Furthermore, cell groups that are sources of major neurotransmission systems in the brain are localized in the brainstem, namely the substantia nigra and ventral tegmental area (VTA) (dopamine), locus coeruleus (noradrenaline), raphe nuclei (serotonin) and reticular formation (acetylcholine) (Nieuwenhuys, 2012). All structures involved in these functions were identified in the present study. It should be noted that the reticular formation and periaqueductal gray were labeled as a single region of interest, given the limitations in spatial resolution and contrast characteristics of our dataset for visualizing individual cell groups.

### Clinical considerations

Alterations of brainstem structure has been and is a current matter of relevance in neurological and neurosurgical clinical practice. Lesions of the brainstem such as embolic or hemorrhagic strokes and tumors can affect the conduit, cranial nerve and integrative functions of the brainstem. Besides the classical neurological syndromes associated with cranial nerve alterations, which are diagnosed using the traditional neurological examination, recently there are a number of other clinical conditions the brainstem is involved in. This is of particular interest given the therapeutic interventions that can be applied using deep brain stimulation (DBS) for neuromodulation. Duret’s work on the involvement of the tegmental reticular formation, thalamic and extra-thalamic pathways as well as of the anterior forebrain circuits with alterations of consciousness, has elucidated the consequences of lesions at the diencephalon-mesencephalic junction leading to coma and prolonged disorders of consciousness (Duret, 1955). Anatomical knowledge of these neuronal loops using neuroimaging to visualize them as targets in current neuromodulation interventions are of clinical relevance given the therapeutic potential of these therapeutic techniques. (Fridman & Schiff, 2014). Interestingly, the cholinergic pedunculopontine tegmental nucleus, which projects to the thalamus and which is involved in conscious behavior, is also involved in gait-balance controls and could be a potential functional target for postural and gait disorders in Parkinson’s disease (Wang et al., 2017). Neurochemical imbalances associated with neurotransmission systems sourcing in brainstem nuclei are also of interest in neuropsychiatric practice and drugs implemented largely in clinical psychiatry affect neurotransmitters. More specifically, selective serotonin reuptake inhibitors (SSRIs) and selective norepinephrine reuptake inhibitors (SNRIs) are routinely used as antidepressants due to their ability to enhance the levels of these neurotransmitters, whereas dopamine antagonists have potent antipsychotic effects by blocking dopamine receptors (Schwartz et al., 2012). Furthermore, cholinergic agents are used in neurodegenerative conditions such as Alzheimer’s disease (Nolte et al., 2016)

### Limitations and future studies

Given the unprecedented quality of the imaging dataset used herein, a cogent way of viewing limitations is by comparing these data with histological observations at microscopic scale. A critical student of neuroanatomy has always as the gold standard a microscopic level of explanation, the cornerstone of traditional neuroanatomy. Neuroimaging is still a few steps away, probably by a factor of ten in spatial resolution, i.e., using approximately 5-micron isotropic voxels, from reaching this level of visualization, although it seems this is rapidly approaching. In this study, we were able to delineate twenty-five nuclear masses and sixteen fiber pathways in the human brainstem using 50-micron isotropic resolution in a postmortem human brainstem. This is still far from the detail provided by microscopic histological examination, where brainstem nuclei and fiber tracts have been identified and delineated using such techniques. It should be noted that in the present study the reticular formation was delineated as a region of interest by exclusion of other structures and guided by histological atlases; the reticular formation per se could not be visualized with certainty using the current methods.

Future studies using combined high-resolution MRI datasets with their own histological sections for comparison (Makris, et al., 2013) should provide further validation and accuracy of anatomical delineations in brainstem anatomy. As MRI technology advances it is expected that higher resolution datasets than the one used in the present study will become available and, thus, histological analyses should be done in these samples as well in a co-registered fashion with the MRI datasets for precise and valid comparisons.

### Conclusions

Using a 50-micron isotropic voxel MRI postmortem dataset of the human brainstem and the 3D Slicer platform set of tools for image analysis, we were able to identify and delineate a large number of nuclear masses, the largest so far to our knowledge, as well as most of the sizeable white matter fiber tracts with conduit and interconnecting nature.

## Acknowledgments

This research was supported in part by R01 MH112748, R01 MH 111917 R21 DA042271 and K24 MH116366 (to Makris N) and NIH grants P41 EB015902, P41 EB015898, R01 CA235589, and HHSN261200800001E (to Kikinis R, Professor of Radiology, Harvard Medical School; Director, Surgical Planning Laboratory, Department of Radiology, and Robert Greenes Distinguished Director of Biomedical Informatics, Brigham and Women’s Hospital; Professor of Medical Image Computing, University of Bremen Institutsleiter, Fraunhofer MEVIS, Bremen).

